## Supplemental Methods, Figures, and Tables for "Bile acids activate the antibacterial T6SS1 in the gut pathogen *Vibrio parahaemolyticus*"

Supplementary Materials and Methods

Supplementary Figures S1-S2

Supplementary Tables S1-S2

Supplementary References

### Supplementary Materials and Methods

**Strains and media:** Bacterial strains used in this study are listed in **Table S1**. *Escherichia coli* strains were grown in 2xYT broth (1.6% [wt/vol] tryptone, 1% [wt/vol] yeast extract, and 0.5% [wt/vol] NaCl) or on Lysogeny broth (LB) agar plates (1% [wt/vol] tryptone, 0.5% [wt/vol] yeast extract, 1% [wt/vol] NaCl, and 1.5% [wt/vol] agar) at 37°C. Media were supplemented with kanamycin (30 µg/mL) or chloramphenicol (10 µg/mL) when appropriate to maintain plasmids. *Vibrio parahaemolyticus* (*Vpara*) strains were routinely grown in MLB (LB containing 3% [wt/vol] NaCl) or on marine minimal media (MMM) agar plates (2% [wt/vol] NaCl, 0.4% [wt/vol] galactose, 5 mM $MgSO_4$, 7 mM $K_2SO_4$, 77 mM $K_2HPO_4$, 35 mM $KH_2PO_4$, 2 mM $NH_4Cl$, and 1.5% [wt/vol] agar) at 30°C. Media were supplemented with kanamycin (250 µg/mL) or chloramphenicol (10 µg/mL) when appropriate to maintain plasmids. For secretion assays, fluorescence reporter assays, and growth assays, *Vpara* were grown in M9 media (1X M9 minimal salts base [FORMEDIUM, MMS0102], 0.8 mM $MgSO_4$, 5 µM $CaCl_2$, 0.4% [wt/vol] glucose, and supplemented with 3% [wt/vol] NaCl).

**Plasmid construction:** Plasmids used in this study are listed in **Table S2**. To construct transcription reporter plasmids, 300 bp upstream of the *vp1400* or 200 bp upstream of the *vpa1270* open reading frames were amplified from the genome of *Vpara* RIMD 2210633 and introduced into the multiple cloning site of pVSV33 (1), upstream of a promoterless *cat* and *gfp* reporter cassette, using the Gibson assembly method (2).

**Construction of deletion strains:** To delete *tfoY* (*vp1028*) in *Vpara* RIMD 2210633, *E. coli* DH5α λ-pir cells containing the pDM4:*tfoY* plasmid were conjugated into *Vpara* RIMD 2210633. Transconjugants were selected on MMM agar supplemented with chloramphenicol. The resulting transconjugants were plated onto MMM agar containing 15% (wt/vol) sucrose for counterselection and loss of the sacB-containing pDM4. Deletions were confirmed by PCR.

**Fluorescence reporter assays and growth assays:** Overnight-grown cultures of *Vpara* strains containing the indicated pVSV33-based plasmids were diluted 1:10 in fresh MLB supplemented with kanamycin to maintain the plasmids, and grown at 30°C for two hours. Then, the cultures were normalized to an $OD_{600}$ = 0.1 in M9 media supplemented with 3% (wt/vol) NaCl and kanamycin, and with either 20 µM phenamil (TOCRIS, 3379; 1 mM stock solution prepared in 20% [vol/vol] DMSO), 20% (vol/vol) DMSO in volumes equivalent to those added for the phenamil treatment, or the indicated final concentrations of sodium deoxycholate (DOC; Sigma, 309710) or sodium taurodeoxycholate hydrate (TDC; Sigma, T0557). The normalized samples were transferred to black, sterile, flat-bottom 96-well microplates (Greiner, 655090) in triplicate (200 µL per well) and grown at 30°C in a BioTek SYNERGY H1 microplate reader with continuous shaking (205 cpm). Every 15 minutes, cell density readings ($OD_{600}$) and fluorescence readings (excitation 479 nm, bandwidth 18 nm; emission 520 nm, bandwidth 18 nm) were taken. GFP reporter activity was analyzed as arbitrary units obtained by dividing the fluorescence reading by the $OD_{600}$ reading after subtracting blank media readings. The experiments were performed at least three times with similar results. Results from a representative experiment are shown.

**Chloramphenicol-resistance reporter assays:** *Vpara* strains containing the indicated pVSV33-based plasmids were streaked onto MLB agar plates supplemented with kanamycin to maintain the plasmids, or kanamycin and chloramphenicol. Plates were incubated overnight at 30°C, and *cat* reporter expression was determined as growth on plates containing chloramphenicol.

**Secretion assays:** *Vpara* strains were grown in MLB at 30°C overnight. Overnight bacterial cultures were diluted 1:10 in fresh MLB and grown at 30°C for two additional hours. Then, bacterial cultures were normalized to $OD_{600}$ = 0.18 in M9 media supplemented with 3% NaCl and incubated for four hours at 30°C in the presence or absence of 20 μM phenamil, 20% (vol/vol) DMSO in volumes equivalent to those added for the phenamil treatment, or 0.025% (wt/vol) DOC. For expression fractions (cells), 0.5 $OD_{600}$ units of cells were harvested and re-suspended in 50 μL of 2X Tris-glycine SDS sample buffer (Novex, Life Sciences) supplemented with 5% (vol/vol) β-mercaptoethanol. Supernatants of volumes equivalent to 5 $OD_{600}$ units were filtered (0.22 μm) and precipitated with deoxycholate and trichloroacetic acid (3). Precipitated proteins were washed twice with ice-cold acetone prior to re-suspension in 20 μL of 100 mM Tris–HCl pH = 8.0, followed by the addition of 20 μL of 2X Tris-glycine SDS sample buffer supplemented with 5% (vol/vol) β-mercaptoethanol. Expression and secretion samples were resolved on TGX stain-free gels (Bio-Rad), transferred onto nitrocellulose membranes, and immunoblotted with custom-made α-VgrG1 (for T6SS1) (4) at a 1:1000 dilution, and α-RpoB (Direct-blot™ HRP anti-*E. coli* RNA Polymerase β antibody; BioLegend, 663907). Results of a representative experiment out of three independent experiments are shown.

### Supplementary Figures

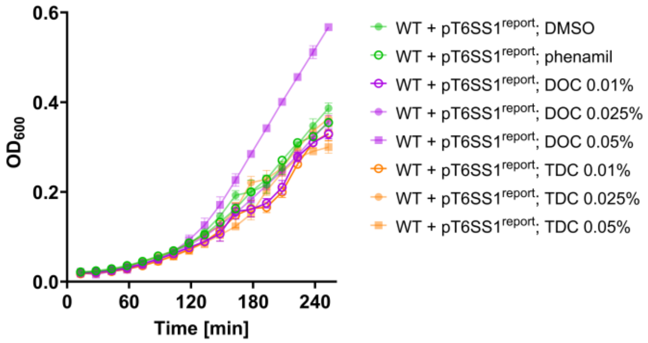


**Fig. S1. Bile acids do not hamper *Vibrio parahaemolyticus* growth.** Growth of *V. parahaemolyticus* RIMD 2210633 carrying a plasmid containing a *gfp* and *cat* reporter cassette fused to the promoter of *vp1400* (pT6SS1$^{report}$), measured as $OD_{600}$. Bacteria were grown at 30°C in M9 media supplemented with 3% (wt/vol) NaCl and kanamycin (250 µg/mL), to maintain the plasmids. Where indicated, the media were supplemented with phenamil (20 µM), DMSO (20% [vol/vol], added to the media as a control at the same volume of the phenamil solution), or the indicated concentrations of DOC or TDC (wt/vol). Data are shown as the mean ± SD, n = 3 independent replicates. The results from a representative experiment out of at least three independent experiments are shown. WT, wild-type.

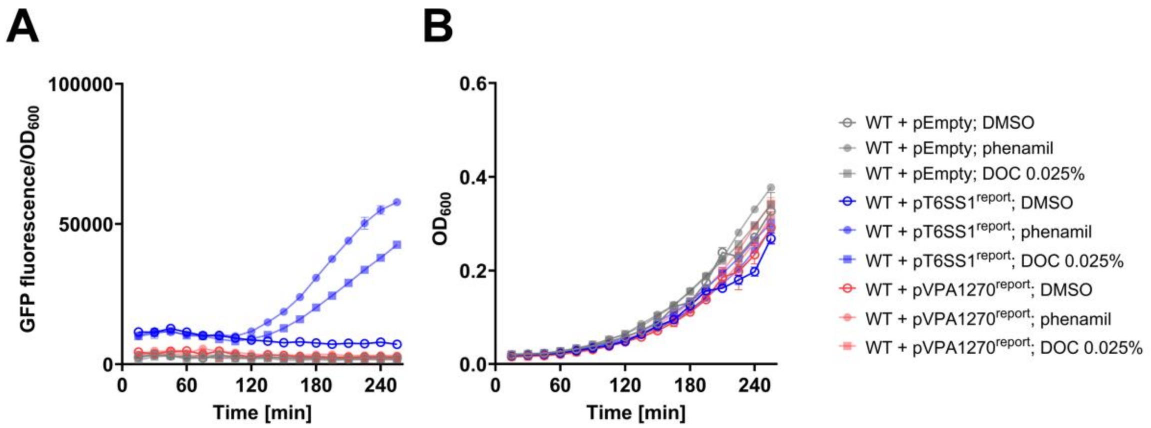


**Fig. S2. Bile acids and phenamil do not have a global effect on transcription and translation in *Vibrio parahaemolyticus*. A)** Fluorescence intensity measured as GFP fluorescence to $OD_{600}$ (arbitrary units), and **B)** growth over time in *V. parahaemolyticus* RIMD 2210633 carrying an empty plasmid (pEmpty), a plasmid containing a *gfp* and *cat* reporter cassette fused to the promoter of *vp1400* (pT6SS1$^{report}$), or a plasmid containing a *gfp* and *cat* reporter cassette fused to the promoter of *vpa1270* (pVPA1270$^{report}$). Bacteria were grown at 30°C in M9 media supplemented with 3% (wt/vol) NaCl and kanamycin (250 µg/mL), to maintain the plasmids. Where indicated, the media were supplemented with phenamil (20 µM), DMSO (20% [vol/vol], added to the media as a control at the same volume of the phenamil solution), or DOC (0.025% [wt/vol]). Data are shown as the mean ± SD, n = 3 independent replicates. The results from a representative experiment out of at least three independent experiments are shown. WT, wild-type.

### Supplementary Tables

**Table S1. A list of bacterial strains used in this study.**

| Strain | Genotype | Comments | Source |
|---|---|---|---|
| *Vibrio parahaemolyticus* RIMD 2210633 | Wild-type | Used for generating deletion strains, secretion assays, and in T6SS1 reporter assays | Obtained from Prof. Kim Orth; (5) |
| *Vibrio parahaemolyticus* RIMD 2210633 Δ*hns* | Δ*vp1133* | Used in T6SS1 reporter assays | (6) |
| *Vibrio parahaemolyticus* RIMD 2210633 Δ*tfoY* | Δ*vp1028* | Used in T6SS1 reporter assays | This study |
| *Vibrio parahaemolyticus* RIMD 2210633 Δ*hcp1* | Δ*vp1393* | Used in secretion assays | (7) |
| *Escherichia coli* DH5α (λ-pir) | K-12 derivative laboratory strain containing λ-pir | Used for plasmid maintenance and cloning | Obtained from Prof. Eric V. Stabb |

**Table S2. A list of plasmids used in this study.**

| Plasmid | Description | Comments | Source |
|---|---|---|---|
| pVSV33 | Kanamycin resistance; promoterless *cat* and *gfp* reporter operon | Parental plasmid for promoter investigation | Obtained from Prof. Eric V. Stabb; (1) |
| pT6SS1$^{report}$ | pVSV33 containing the 300 bp upstream of the *vp1400* start codon transcriptionally fused to the GFP and chloramphenicol-resistance reporter cassette | Used to investigate expression from the *vp1400* promoter of *Vibrio parahaemolyticus* RIMD 2210633 | This study |
| pVPA1270$^{report}$ | pVSV33 containing the 200 bp upstream of the *vpa1270* start codon transcriptionally fused to the GFP and chloramphenicol-resistance reporter cassette | Used to investigate expression from the *vpa1270* promoter of *Vibrio parahaemolyticus* RIMD 2210633 | This study |
| pDM4:*tfoY* | a $Cm^R$ and $ori_{R6K}$-containing suicide vector harboring 1 kb upstream and 1 kb downstream of *vp1028* in its MCS | Used to delete *tfoY* in *Vibrio parahaemolyticus* RIMD 2210633 | (8) |
